# V1 superficial layers create a saliency map that feeds forward to the parietal cortex for attentional guidance

**DOI:** 10.1101/2025.04.10.648136

**Authors:** Chen Liu, Chengwen Liu, Laurentius Huber, Li Zhaoping, Peng Zhang

**Affiliations:** State Key Laboratory of Brain and Cognitive Science, Institute of Biophysics, Chinese Academy of Sciences, Beijing 100101, China; School of Life Sciences, University of Chinese Academy of Sciences, Beijing 100049, China; Cognition and Human Behavior Key Laboratory of Hunan Province, School of Educational Science, Hunan Normal University, Changsha, China; Functional MRI Core, National Institute of Mental Health, National Institutes of Health, Bethesda, MD, United States; University of Tübingen, Max Planck Institute for Biological Cybernetics, Tübingen, Germany

**Keywords:** Saliency, Attention, Laminar fMRI, VASO CBV

## Abstract

A salient visual object with a distinct feature from the surrounding environment automatically captures attention. While the saliency signals have been found in many brain regions, their source remains highly controversial. Here, we investigated the neural origin of visual saliency using cortical layer-dependent functional magnetic resonance imaging (fMRI) of cerebral blood volume (CBV) at 7 Tesla. Behaviorally, human observers were better at detecting foreground bars with a larger orientation contrast from uniformly oriented background bars. Saliency-sensitive signals were strongest in the superficial layers of the primary visual cortex (V1), and in the middle layers of the intraparietal sulcus (IPS) of the parietal cortex. Layer-dependent effective connectivity revealed the transmission of saliency signals along the feedforward pathway from V1 to IPS. Furthermore, behavioral sensitivity to the foreground stimulus correlated significantly with the fMRI response in the superficial layers of V1. Our findings provide mesoscale evidence that a visual saliency map is created by iso-feature suppression through lateral inhibition in the superficial layers of V1, and then feeds forward to attentional control brain regions to guide attention and eye movements.

## Introduction

Only a small fraction of sensory inputs can be selected by our attention for further processing. Attention is guided both endogenously by top-down factors and exogenously by bottom-up factors (Baluch & Itti, 2011; Bisley & Goldberg, 2010; Buschman & Kastner, 2015; Hopfinger et al., 2000; Moore & Zirnsak, 2017). Although the bottom-up guidance is simpler, it remains controversial which brain area computes saliency from exogenous visual input. It is believed that the brain generates a saliency map from feature contrast to guide attention shifts and eye movements (Itti et al., 1998; Koch & Ullman, 1985; Li, 2002; Treisman & Gelade, 1980). According to a traditional and dominant computational model (Itti & Koch, 2000, 2001; Koch & Ullman, 1985), different feature maps (e.g., color, luminance, orientation, etc.) are generated by center-surround mechanisms and then combined into a feature-agnostic master map of saliency, presumably in a high-order brain region such as the parietal cortex. However, it has been shown that saliency behavior can also be explained by the V1 Saliency Hypothesis (V1SH) (Li, 2002, 2008), according to which the saliency map is generated by intracortical mechanisms in the primary visual cortex (V1) and represented by the firing rates of feature-selective neurons.

Visual saliency-related signals have been observed in the frontal (Moore & Armstrong, 2003; Thompson & Bichot, 2005) and parietal cortices (Bisley & Goldberg, 2010; Chen et al., 2020), V4 (Mazer & Gallant, 2003) and V1 (Yan et al., 2018; Zhang et al., 2012), as well as subcortical regions including the pulvinar of the thalamus (Shipp, 2004) and the superior colliculus (SC) of the midbrain (Fecteau & Munoz, 2006; White, Berg, et al., 2017; White, Kan, et al., 2017). An important question is which area generates the saliency signals originally and which areas merely receive and utilize the saliency signals generated in another area. For example, a recent study showed that inactivation of the posterior parietal cortex in monkeys significantly weakened the relationship between image salience and visually guided behavior (Chen et al., 2020), this could suggest that the parietal cortex is the original source of the saliency signals, but this is also consistent with the idea that parietal cortex utilizes the saliency signals and combine them with top-down signals for guiding behavior (Bisley & Goldberg, 2010). Using salient but subliminal visual inputs, a human neuroimaging study found responses to saliency in the earliest (C1) component of event-related potentials (ERPs), as well as fMRI responses to saliency in V1 but not in the parietal cortex (Zhang et al., 2012). One monkey electrophysiological study reported that the latency of the saliency signals in SC is shorter than that in V1 (White, Kan, et al., 2017), but this short SC latency is longer than the latency of the saliency signals in V1 reported in another monkey study (Yan et al., 2018). We aim to shed light on the origin of saliency signals through a high-resolution fMRI study, which allows us to examine laminar patterns of responses to saliency signals. As will be explained later, the laminar patterns are diagnostic of the sources of the saliency signals, and we will focus on the question of whether the source for the saliency signals is in V1, IPS, or SC.

Recent advances in high-resolution fMRI at ultra-high magnetic field have enabled non-invasive imaging of mesoscale functional units, such as cortical layers and columns, in the human brain at submillimeter resolution (Cheng, 2018; Huber et al., 2020; Vizioli et al., 2023). Using submillimeter blood oxygen level dependent (BOLD) fMRI techniques, feedforward and feedback pathways of stimulus-driven activity and top-down attentional modulation have been investigated in the human visual cortex (de Hollander et al., 2021; Lawrence et al., 2019; Liu et al., 2021; Van Mourik et al., 2023). However, the laminar specificity of T2* weighted BOLD fMRI is limited by the draining vein effect (Fukuda et al., 2021; Huber et al., 2019; Kay et al., 2019; Uludag & Havlicek, 2021). The Vascular Space Occupancy (VASO)-based cerebral blood volume (CBV) technique is an inversion recovery MRI technique that detects changes in blood volume by nulling the signal from blood water (Lu et al., 2003, 2004), particularly in small vessels such as arterioles and capillaries. Therefore, compared to T2*w BOLD signals that are sensitive to blood oxygen in large veins, CBV changes are more closely related to the site of neuronal activation, offering higher spatial specificity to resolve layer-dependent signals across cortical depth, despite having relatively lower functional sensitivity (Finn et al., 2019; Huber et al., 2017).

**Figure 1.**
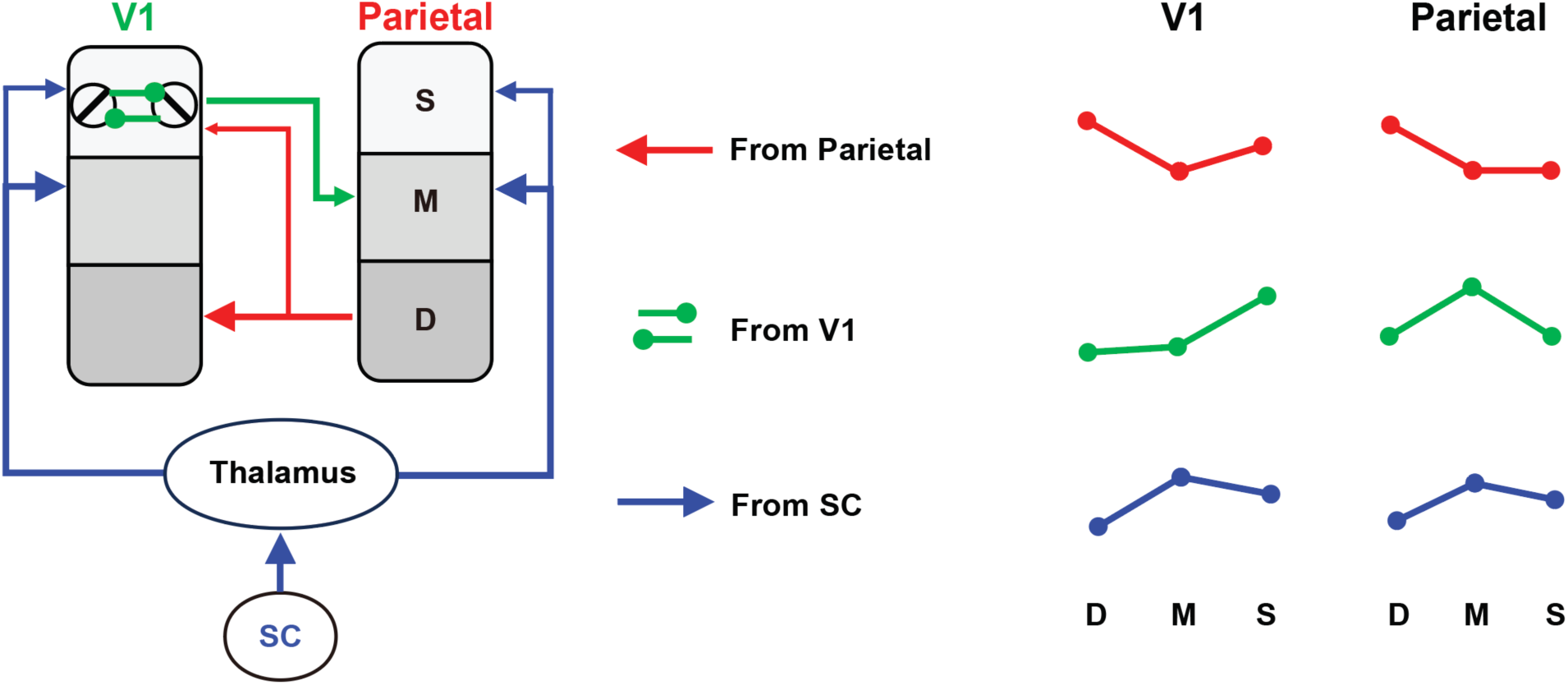
Three possible hypotheses on the origins of saliency signals and their consequent laminar patterns of the saliency signals through feedforward and feedback neural pathways. In the feature combination hypothesis, saliency signal initially emerged in frontoparietal areas and then modulate neural activity of parietal and early visual cortices through feedback connections (Red lines); In V1SH, saliency signals initially emerge via intracortical interactions mediated by the horizontal connections in superficial layers of V1 (primary visual cortex). These signals can then feedforward to middle layers of parietal cortex (Green lines); In subcortical computation hypothesis, saliency signals are generated in SC (superior colliculus) and then modulate middle and superficial layer activity of cerebral cortex through thalamo-cortical pathways. On the right, relative strengths of saliency signals in the deep (D), middle (M), and superficial (S) layers of V1 and parietal cortex that should arise from each hypothesis are shown.

Different brain areas for creating the saliency signals should lead to different laminar patterns of the distributions of these saliency signals in themselves as well as in other brain areas as the saliency signals are propagated through feedforward and feedback connections. One can therefore examine these laminar patterns by laminar fMRI techniques to diagnose the source of the saliency signals. Figure 1 schematizes the laminar pattern of saliency-related signals in V1 and parietal cortex according to each of the three possible origins of the saliency signal: parietal cortex, V1, and SC, based on the canonical neural circuits in the primate visual system (Felleman & Van Essen, 1991). If the parietal cortex generates the saliency map and modulates V1 activity through feedback connections, the laminar pattern of saliency-related signals should be consistent with cortico-cortical feedback, stronger in the deep and superficial layers of V1 and in the deep layer of the parietal cortex. According to the V1 Saliency Hypothesis (V1SH), saliency signals emerge through intracortical interactions mediated by horizontal connections in the superficial layers of V1. Consequently, the strongest saliency-related activity should be observed in the superficial layers of V1 and in the middle layers of high-level cortex that receive feedforward input. Finally, if the saliency signals originate in the SC, they would modulate activity more strongly in the middle and superficial cortical layers through the tecto-thalamo-cortical pathway (Shipp, 2003, 2004, 2007).

In the current study, to reveal the neural source of visual saliency, we employed submillimeter fMRI at 7 Tesla with a CBV-weighted VASO fMRI technique to investigate cortical depth-dependent responses to a foreground stimulus consisting of four bars presented at different orientation contrasts (θ = 90°, 15° or 0°) relative to a background of iso-oriented bars (Figure 2). The laminar profile of the saliency-sensitive fMRI response, its correlation with behavioral sensitivity to saliency, and the layer-dependent functional connectivity collectively support the hypothesis that the saliency map is generated in the superficial layers of V1 and subsequently projects to attention-control brain regions, such as the parietal cortex.

## Results

### Behavioral sensitivity was higher to a foreground region with a larger orientation contrast against the background

In a psychophysical experiment, the bar stimulus was presented for 200 ms (Figure 2A). The foreground bars were presented either to the lower-left or the lower-right of fixation, at an eccentricity of 4.3°, and participants pressed one of two buttons to report whether the foreground bars were at lower-left or lower right. The orientation of the uniformly oriented background bars was randomly chosen for each trial, while the foreground bars had an orientation contrast θ of either 90° or 15° relative to the background bars. Across trials, the Michaelson luminance contrast between the bars (all of which had the same luminance) and the uniform gray background was adaptively adjusted using a 3-down-1-up staircase procedure, implemented independently for the two θ conditions, to identify a threshold contrast *C_threshold_* (at which the participant can perform the task at 80% accuracy. The results showed that luminance contrast sensitivity *S_behavior_* = 1/*C_threshold_* was significantly higher for a foreground with a larger orientation contrast θ (Figure 2B, 90° vs. 15°, t_19_ = 13.829, p < 0.001). Specifically, the behavioral sensitivity to detect the 90° foreground (*S_behavior_*(90°) = 18.719) was approximately twice that of the sensitivity to detect the 15° foreground (*S_behavior_*(15°) = 9.720) (Figure 2B). It should be noted that, at our luminance contrast sensitivity *S_behavior_* to detect the foreground, the luminance of all the bars were presented at a suprathreshold level. Hence, the luminance contrast threshold *C_threshold_* measured in our task does not reflect the visibility of the entire stimulus but rather the saliency of the foreground bars in the context of the background texture. When the orientation contrast θ or saliency is higher, the foreground is more easily detected and localized, so that our participants could do our task at a lower luminance contrast of the bars. Hence, our luminance contrast sensitivity *S_behavior_* is an assessment of the saliency of the foreground bars, and we also call *S_behavior_* our *behavioral saliency score*. Furthermore, to correlate with the fMRI signals for saliency later, we define

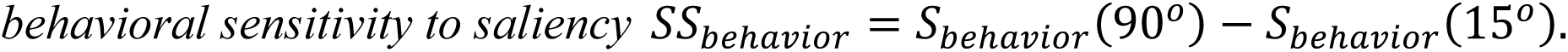

**Figure 2.**
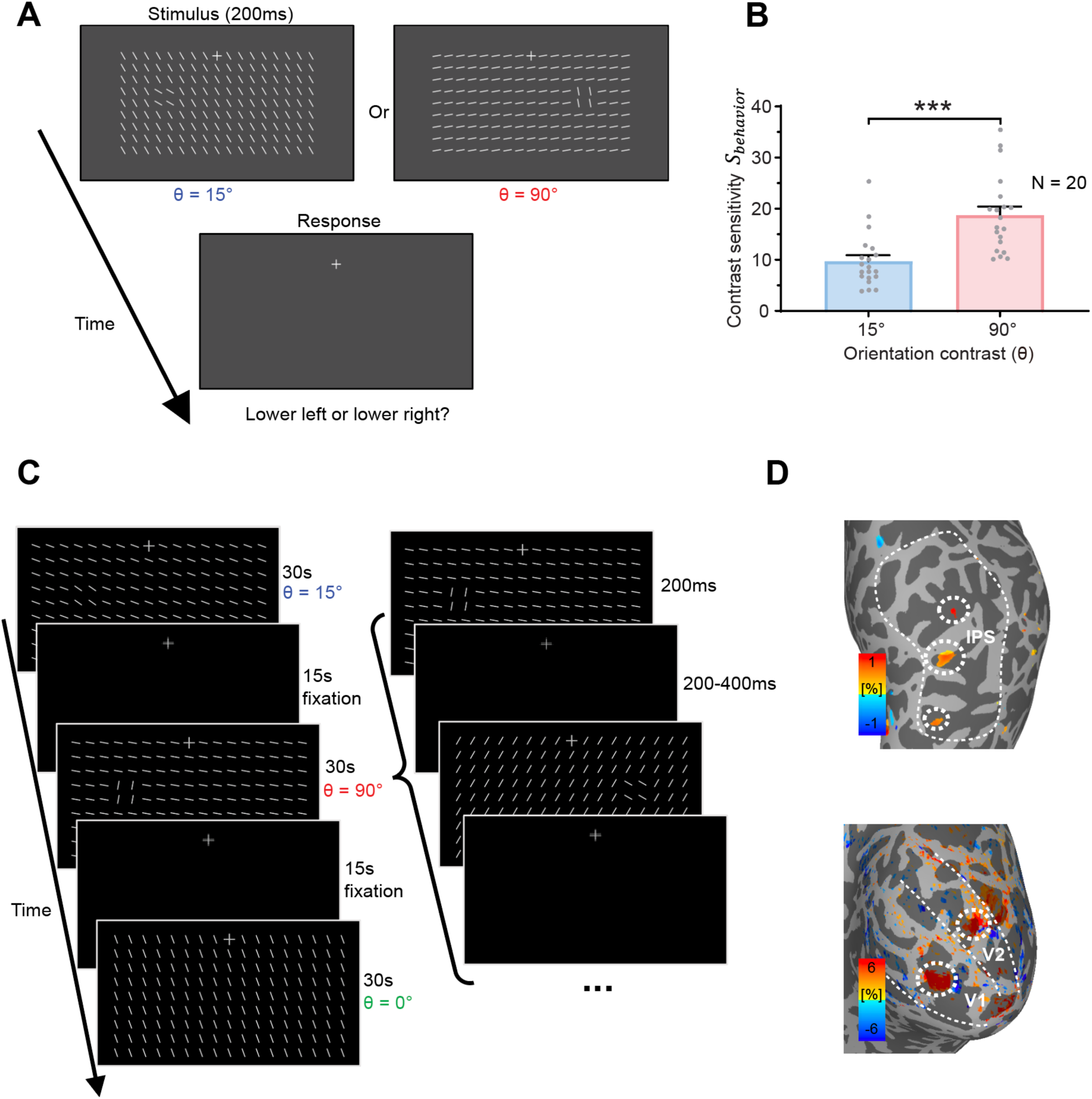
(A) Stimulus and procedure of the psychophysical experiment. The orientation contrast between foreground and background bars is defined as θ. (B) Luminance contrast sensitivity (*S_behavior)_* for detecting the 90° and 15° foreground. Each dot represents data from one participant. Error bars represent standard errors of the mean (SEM). *** p < 0.001. (C) Stimulus and procedure of the fMRI experiment. Texture stimuli with three θ conditions were presented in separate blocks. The foreground bars were randomly presented to the lower-left or lower-right of fixation in each block. (D) Localizer activations corresponding to the retinotopic location of the foreground bars in V1 and V2 of the early visual cortex and IPS of the parietal cortex (thresholded at p < 0.05 uncorrected in a representative participant), as indicated by the dashed circles. Dashed lines mark the borders of the brain areas.

### Saliency-sensitive responses were strongest in the superficial layers of V1 and in the middle layers of IPS

In the 7T fMRI experiment, a slice-saturation slab-inversion VASO (SS-SI-VASO) sequence with 0.82-mm isotropic resolution was used to measure CBV-weighted signals across cortical depth from the posterior part of the brain (Huber et al., 2014, 2017). In each inversion TR, blood-nulled and not-nulled 3D-EPI volumes were acquired to minimize T2*-weighted BOLD contamination. The imaging slab was oriented in an oblique-coronal orientation to cover the early visual cortices and the parietal cortex from the dorsal part of the occipito-parietal lobe (Figure S1C). According to the literature and hypotheses on the neural origin of visual saliency (Figure 1), our analyses focused on V1 and V2 of the early visual cortex, and the intraparietal sulcus (IPS) of the parietal cortex.

The stimulus and procedure of the fMRI experiment are illustrated in Figure 2C. Bar stimuli with three orientation contrast conditions (θ = 90°, 15° and 0°, randomly selected to be either clockwise or counterclockwise to the orientation of the background bars) were presented in the lower visual field in separate 30-second blocks, interleaved with 15-second fixation period. Within a stimulus block, θ is fixed, and each stimulus was presented for 200 ms, with a random inter-stimulus interval (ISI) ranging from 200 to 400 ms. In each stimulus, the iso-oriented background bars had a random orientation, and the foreground bars appeared randomly either to the lower left or lower right of fixation. During the experiment, participants were instructed to maintain fixation and to detect small luminance changes of the fixation cross. In two localizer runs, naturalistic stimuli of the same size and location as the foreground bars were presented in the lower-left and lower-right visual fields in separate blocks (Figure S2). The cortical ROIs of the foreground bars are shown in Figure 2D for a representative participant.

**Figure 3.**
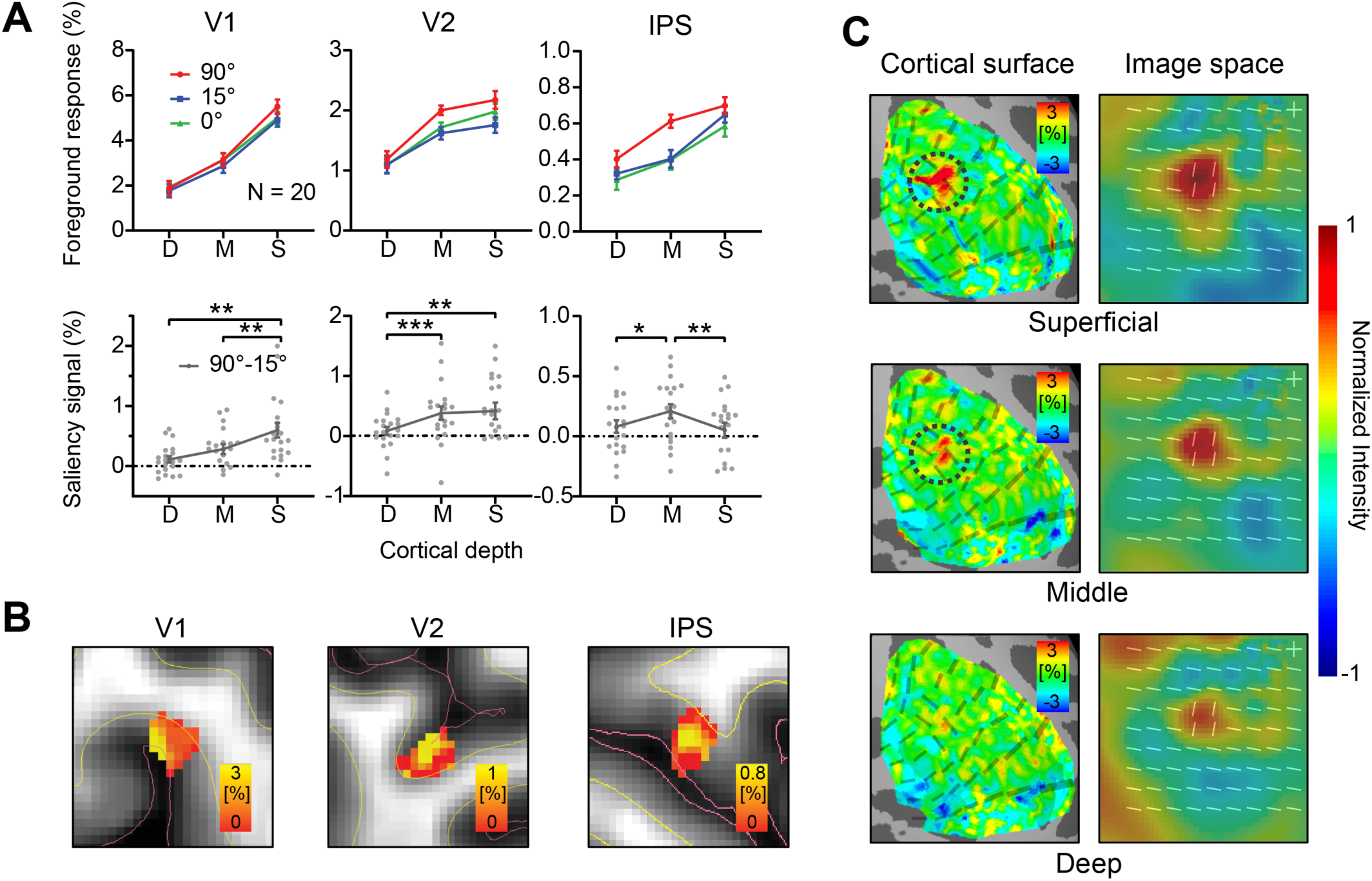
(A) Upper: Normalized CBV responses (*S_fMRI_* in percent signal change) to the orientation foregrounds in V1, V2, and IPS. Lower: CBV response difference between the 90° and 15° foreground conditions 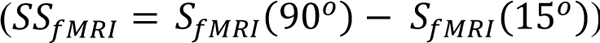. Each gray dot represents data from one participant. Error bars represent SEM. *, ** and *** indicate p < 0.05, p < 0.01 and p < 0.001, respectively. D: deep, M: middle, S: superficial. (B) Volume activation maps of the saliency-sensitive response *SS_fMRI_* in the foreground ROIs of V1, V2, and IPS for a representative participant. Red lines indicate the boundary between gray matter (GM) and cerebrospinal fluid (CSF). Yellow lines indicate the boundary between GM and white matter (WM). (C) Left: Surface activation maps of saliency-sensitive responss (*SS_fMRI_*, in percent signal change) in different cortical depths of V1 in a representative participant (same participant as in B). Dashed circles indicate the location of foreground on the cortical surface. Right: Saliency maps (averaged across all participants) in image space.

To quantify our results and to account for individual differences in mean response amplitude, we define the normalized CBV foreground response as 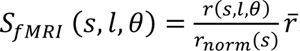. In which, *r*(*s*, *l*, *θ*) is the original CBV response for subject *s*, layer *l*, at foreground orientation contrast *θ*, 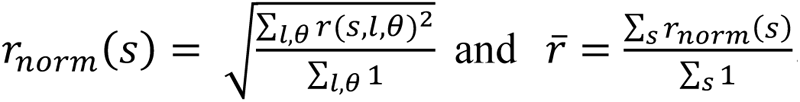. CBV-weighted fMRI responses in the foreground ROIs *S_fMRI_* in V1, V2 and IPS are plotted as a function of cortical depth in Figure 3A (see Figure S3 for the BOLD response profiles).

A two-way repeated measures (rm) ANOVA with cortical depth (deep, middle, and superficial) and orientation contrast (θ = 90°, 15°, and 0°) as within-subject factors was performed on the normalized foreground response *S_fMRI._*. A significant effect of orientation contrast θ was found in all three brain regions (V1: F_2, 38_ = 7.080, p = 0.002; V2: F_2, 38_ = 4.763, p = 0.014; IPS: F_2, 38_ = 4.673, p = 0.015), indicating stronger responses to the foreground with larger orientation contrast (i.e., 90° compared to 15° and 0° conditions). However, CBV responses to the 15° foreground were slightly reduced compared to the 0° condition. This is due to the partial volume effect of background suppression (Figure S4). Similar effects of background suppression were found for the 90° and 15° conditions. Therefore, to avoid the partial volume effect of background suppression, *the saliency-sensitive fMRI response* 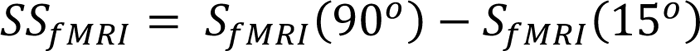 was calculated as the difference between the fMRI responses to the 90° foreground and the 15° foreground conditions (Figure 3A, lower).

The amplitude of *SS_fMRI_* differed significantly across cortical depth in V1, V2, and IPS (V1: F_2, 38_ = 9.696, p = 0.002; V2: F_2, 38_ = 9.833, p = 0.001; IPS: F_2, 38_ = 5.421, p = 0.008). In V1, the saliency-sensitive response was strongest in the superficial layers (S vs. D: t_19_ = 3.408, p = 0.003; S vs. M: t_19_ = 3.660, p = 0.002). In V2, it was stronger in the superficial and middle layers than in the deep layers (S vs. D: t_19_ = 3.232, p = 0.004; M vs. D: t_19_ = 4.582, p < 0.001). In IPS, *SS_fMRI_* was strongest in the middle layers (M vs. D: t_19_ = 2.659, p = 0.015; M vs. S: t_19_ = 2.983, p = 0.008). Saliency-sensitive responses calculated by the original CBV responses showed a similar pattern of results (Figure S5). The laminar profiles of the response difference between the 90° and 0° conditions, *S_fMRI_*(90°) - *S_fMRI_*(0°), were also qualitatively similar (Figure S6). In Figure 3B, the activation map in a representative participant clearly showed the strongest saliency sensitivity *SS_fMRI_* in the superficial layers of V1, in the middle and superficial layers of V2, and in the middle layers of IPS, consistent with the laminar profiles for a V1 origin of saliency in Figure 3A. To better illustrate the V1 saliency map across cortical depths, saliency-sensitive responses *SS_fMRI_* were shown on the cortical surface in the right hemisphere of the representative subject (Figure 3C, left). The saliency maps in the image space were reconstructed using a population receptive field (pRF) model, shown in the right panels of Figure 3C. Saliency signals were strongest in the superficial layer of V1 at the location of foreground bars. These findings are consistent with the V1 hypothesis in Figure 1, in which the saliency signals initially emerge by horizontal interactions in the superficial layers of V1 and subsequently feed forward to the middle layers of IPS.

### Behavioral sensitivity to saliency correlated with the saliency-sensitive fMRI response in the superficial layers of V1

To investigate whether the fMRI response can predict the behavioral sensitivity to the foreground bars, we calculated Pearson’s correlations between the behavioral sensitivity to saliency *SS_behavior_* and the saliency-sensitive fMRI response *SS_fMRI_*. For each brain region, the cortical depth with the strongest saliency-sensitive fMRI response was selected. *SS_behavior_* showed a significant correlation with *SS_fMRI_* in the superficial layers of V1 (Figure 4A, r = 0.542, p = 0.014), but not with those in the superficial layers of V2 (r = -0.218, p = 0.356) or the middle layers of IPS (r = -0.203, p = 0.390). The correlation in V1 remained significant after family-wise error (FWE) correction by permutation test (Figure 4B). The correlation results supported that the saliency-sensitive fMRI response in the superficial layers of V1 can predict the behavioral sensitivity to saliency.

**Figure 4.**
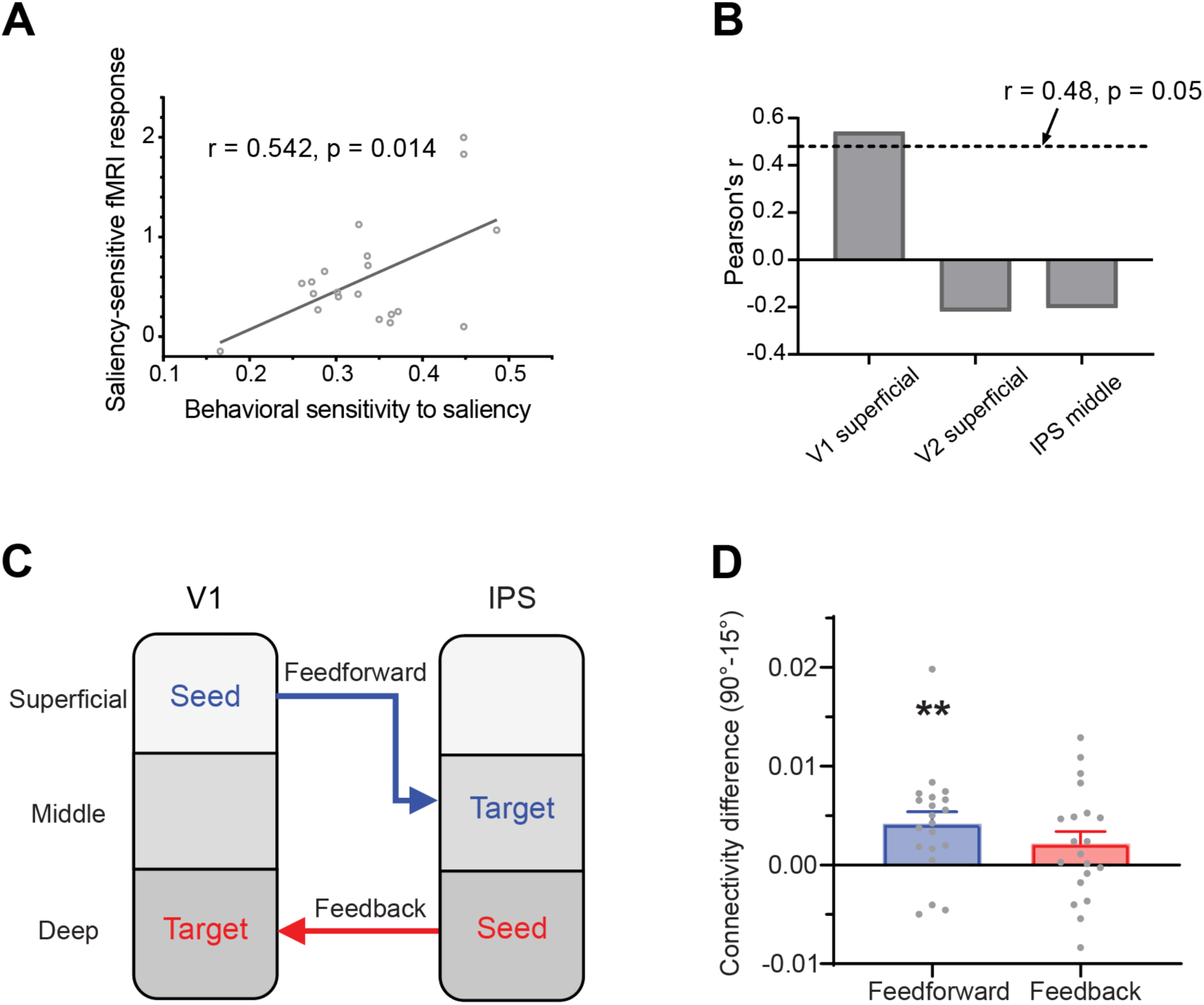
(A) Behavioral sensitivity to saliency *S_behavior_* i.e., the difference in luminance contrast sensitivity to detect the two different foregrounds, is correlated with fMRI saliency-sensitive response *SS_fMRI_*, in the superficial layers of V1, giving a significant Pearson’s correlation coefficient r. Each circle represents one participant. (B) Pearson’s correlation coefficients r in the superficial layers of V1 and V2, and in the middle layer of IPS. The dashed line indicates the significance threshold after family-wise error (FWE) correction by a permutation test. (C) The feedforward and feedback pathways of saliency signals between V1 and IPS, assessed by the layer-specific generalized psychophysiological interaction (gPPI) analysis of functional connectivity. (D) Functional connectivity difference (90° vs. 15°) of the two pathways in C. Each gray dot represents a single participant. Error bars represent the standard errors of the mean across participants. ** indicates p < 0.01.

### Layer-dependent effective connectivity revealed saliency transmission from V1 to IPS

A generalized psychophysiological interaction (gPPI) analysis was conducted to examine layer-dependent effective connectivity of saliency signals between V1 and IPS. For the feedforward connection, the superficial layer of V1 was used as the seed ROI, and the middle layer of IPS served as the target ROI. For the feedback connection, the seed ROI was the deep layer of IPS, and the target ROI was the deep layer of V1 (Figure 4C). The gPPI results revealed a significant connectivity difference between the 90° and 15° foreground conditions for the feedforward connection from V1 superficial to IPS middle (t_19_ = 3.407, p = 0.006, Holm corrected). In contrast, no significant difference was observed for the feedback connection from IPS deep to V1 deep (t_19_ = 1.719, p = 0.102) (Figure 4D). Due to the low signal-to-noise ratio of the VASO signal, here we used the BOLD signals for the gPPI analysis. The gPPI results using VASO and BOLD signals were similar (Figure S7), both showing a significant feedforward connection from V1 to IPS. The connections across all cortical layers are shown in Figure S7, indicating that the middle layer of IPS received significant feedforward input from V1. The layer-dependent connectivity results further support the hypothesis that the saliency signal propagates along the feedforward pathway from V1 to IPS.

### Fixation stability was not affected by the presence of foreground bars

Since participants were performing a fixation task in the fMRI experiment, the fixation stability was unlikely to be affected by the presence of the foreground. To confirm that participants could maintain fixation, we monitored the eye movements of 12 participants in a control experiment using the same stimuli and fixation task as in the fMRI experiment. Fixation stability was assessed using the bivariate contour ellipse area (BCEA) of 95% fixations (Steinman, 1965). BCEA results showed no significant difference among the three θ conditions (F_2, 22_ = 0.291, p = 0.750, BF_10_ = 0.227) (Figure S8). Therefore, our findings were unlikely to be due to unstable fixation.

## Discussion

The neural source of bottom-up saliency has been highly controversial. Using 7T VASO-CBV fMRI at submillimeter resolution, we investigated layer-dependent responses to salient visual input in the early visual cortex and the parietal cortex of human participants, and the functional connectivity between these brain regions to elucidate how saliency signals are propagated. Saliency-sensitive responses were strongest in the superficial layers of V1, and in the middle layers of IPS. Meanwhile, behavioral sensitivity to saliency correlated significantly with the fMRI saliency-sensitive responses in the superficial layers of V1, but not with those in the middle layers of IPS. Furthermore, layer-dependent effective connectivity revealed significant feedforward transmission of saliency signals from V1 to IPS. These findings support the hypothesis that a bottom-up saliency map is created by contextual influences mediated by horizontal connections, which are most abundant in the superficial layers of V1, and is subsequently propagated to other brain regions, such as IPS and SC, to guide attention and eye movements.

While the V1 saliency hypothesis (V1SH) is supported by substantial behavioral evidence (Koene & Zhaoping, 2007; Li, 2008; Zhaoping, 2012; Zhaoping & May, 2007; Zhaoping & Zhe, 2015), only a few neuroscience studies have directly tested it. Using EEG and fMRI, (Zhang et al., 2012) studied the neural responses to salient foregrounds resembling our stimuli, but the bars were made subliminal by backward masking so that observers were at chance level in identifying the foreground location, while the saliency effect of the foreground (manifested as its cueing effect in discriminating a subsequent probe) remained evident. A more salient foreground (by a larger orientation contrast) evoked a stronger C1 component of occipital ERPs, and 3T fMRI results revealed saliency signals in V1 but not in IPS. Although these findings provide strong neuroimaging support for the V1SH, several limitations should be noted. First of all, top-down attention may not be completely abolished by backward masking, since it can modulate stimulus processing even without awareness of visual stimuli (Bahrami et al., 2007; Kanai et al., 2006). In addition, C1 amplitude can also be modulated by top-down attention (Kelly et al., 2008; Rauss et al., 2009, 2012). Finally, the lack of a saliency signal in IPS could be due to the limited spatial resolution of 3T fMRI to isolate the retinotopic signal of the foreground. Meanwhile, for supraliminal salient visual inputs, saliency-sensitive responses in parietal cortex have been observed in fMRI and neurophysiological studies (Bisley & Goldberg, 2010; Geng & Mangun, 2009; Gottlieb et al., 1998). In one neurophysiological study (Yan et al., 2018), neural activity in monkey V1 showed both early (40-60 ms after stimulus onset) and late (80-200 ms) responses to a salient orientation singleton bar in a background of uniformly oriented bars, with the early component being present even during the fixation task, prior to any experience in the singleton detection task. However, another study (White, Kan, et al., 2017) found that saliency-related signals appeared earlier in the SC (around 65 ms) than in V1 (around 121 ms). Therefore, neuroscience studies have found saliency-related signals in multiple brain regions, and it remains controversial as to which brain regions are, respectively, the sources and recipients of the saliency signals.

With a large brain coverage and a submillimeter resolution, the laminar fMRI results in our study provide important mesoscale evidence for the cortical microcircuitry underlying the saliency computations for bottom-up attention in humans. The laminar response patterns from multiple brain regions (Figure 3A/B) and the layer-dependent connectivity (Figure 4C/D) are consistent with a canonical feedforward pathway from V1 to IPS (Felleman & Van Essen, 1991; Markov et al., 2014). Since the intracortical horizontal connections in V1 are most abundant in the superficial layers (Gilbert & Wiesel, 1983; Rockland & Lund, 1983; Angelucci et al., 2017), and since contextual influences mediated by these connections is a key mechanism underlying the saliency computation in the V1 model (Li, 2002), our finding that the strongest saliency-related signal is in the superficial layers of V1 strengthens the evidence for V1 as the source of the saliency signals. While the previous studies cannot rule out the superior colliculus (SC) as a potential source for saliency computation, our findings provide evidence against this possibility. If the saliency map was generated in the SC, the saliency signals should be stronger in the middle layer than in the superficial layer of V1 (Figure 1), since the thalamic input mediated from the SC via the tecto-thalamo-cortical pathway should be strongest in the middle layer of V1 (Shipp, 2003, 2007). Our laminar results (Figure 3) are not consistent with this prediction. Instead, they are aligned with recent evidence that visual responses in the SC of awake primates depend on signals routed through the LGN and V1 (Katz et al., 2024): deactivation of the LGN abolished SC responses to visual stimuli. It is therefore more likely that the saliency signals are generated in V1 and then project to the superficial layers of the SC to guide attention and eye movements (May, 2006; Shipp, 2004).

In contrast to the results reported by (Zhang et al., 2012), we observed significant saliency-sensitive signals in IPS (Figure 3A/B). This discrepancy may be attributed to the higher spatial resolution of our 7T fMRI data, which allowed better isolation of fMRI responses to the foreground from those to the background bars. Another possible reason is the visibility of the texture stimuli, which were made invisible by backward masking in Zhang et al. (2012). However, given the low-level mechanisms in V1 and the feedforward transmission of saliency signals, it is unlikely to be strongly influenced by conscious awareness (He et al., 1996; He & MacLeod, 2001; Rees et al., 2002; Tong, 2003). By making the stimuli invisible, the previous study largely eliminated conscious awareness-related feedback from a high-level cortex such as the IPS, while the saliency signals were still observed in V1. However, as a consequence, the feedforward propagation of the saliency signals from V1 to IPS cannot be illustrated due to the lack of signal in the high-level region. Overcoming the limitations of the previous study, our study represents important advances in demonstrating the neural origin and recipient of saliency signals. We reveal the V1 origin and feedforward propagation of saliency signals from V1 to IPS, by allowing the manifestation of the saliency signals in the parietal cortex and by layer-specific response and connectivity analyses. Furthermore, the laminar response pattern for horizontal contextual influences in the superficial layers of V1 provides cortical support for the proposed mechanism of saliency computation in the V1 model.

To summarize, mesoscale evidence at the level of cortical layers resolves key discrepancies in the neural origin of saliency signals. Our results demonstrate that visual saliency is initially created by iso-feature suppression in the superficial layers of V1 and then propagates to higher-order attentional control brain regions to guide attention and eye movements. These findings not only shed light on the neural and computational mechanisms of visual attention, but also provide valuable guidance for the development of biologically inspired artificial visual systems.

## Methods and Materials

### Participants

Twenty healthy volunteers (9 females, aged 22-42 years) participated in the psychophysical and fMRI experiments. The sample size was determined based on the large effect (Cohen’s f > 0.4) of orientation contrast on V1 responses in a previous study (Zhang et al., 2012), as calculated by G*Power (Faul et al., 2009). A sample size of 20 participants was sufficient to achieve 90% statistical power for detecting a large effect in a within-subjects ANOVA. Twelve volunteers (5 females, 22-30 years of age) participated in the control movement with eye movement recordings, 6 of them also participated in the fMRI experiment. All participants had normal or corrected-to-normal vision and gave written informed consent before the experiments. Experimental protocols were approved by the Institutional Review Board of the Institute of Biophysics, Chinese Academy of Sciences (No. 2012-IBP-011).

### Stimuli and procedures

Visual stimuli were generated in MATLAB (Mathworks Inc.) with psychophysics toolbox extension (Brainard, 1997; Pelli, 1997) on a Windows 10 operating system computer. Stimuli and procedures were illustrated in Figure 2A&2C. Each texture stimulus (12.5 by 6.5 degrees of visual angle) consisted of 17 × 9 bars of 0.5° × 0.05° in visual angle presented in the lower visual field on a gray background (*L*_0_ 43.8 cd/m^2^), with 0.75° spacing between the bars. In the foreground present conditions, all bars were identical except for a foreground of 2 × 2 bars titled either 90° or 15° (orientation contrast θ) relative to the background bars. The foreground bars were located at 4.3° of eccentricity either to the lower-left or lower-right of the fixation. In the uniform texture condition, all bars were uniformly oriented. The orientation of background bars was randomly chosen from 0° to 180°.

#### Psychophysical experiment

Visual stimuli were presented on a Cambridge Research System Display++ LCD monitor (32 inches, 1920 × 1080 pixels, 120-Hz refresh rate) at a viewing distance of 100 cm. The head of subject was stabilized using a chin rest.

A bright fixation cross (73 cd/m^2^) was shown at the center of the screen. Participants were informed to keep fixation throughout the experiment. A trial began after a button press. After 1 s of fixation, the texture stimulus was presented for 200 ms, followed by another 1 s of fixation for the participant to respond (Figure 2A) before the next trial began. Participants were instructed to make 2-alternatives-forced-choice response by pressing one of two buttons to indicate whether the foreground appeared in the lower left or lower right visual field. Bar luminance *L_bar_* (ranged from 43.8 to 78.9 cd/m^2^) was adjusted by a 3-down-1-up staircase procedure to determine the participant’s contrast sensitivity for detecting the foreground. The luminance contrast of the bars from the uniform gray background was defined by the Michaelson contrast 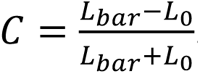. Two blocks of trials (120 trials each) were collected for each orientation contrast condition. Each block consisted of two independent staircases (60 trials each) that were randomly intermixed across trials. Thus, a total of four 60-trial staircases were collected for each condition. A psychometric curve was fitted by a Weibull function relating the performance in each condition to the contrast of the bars. The contrast detection threshold *C_threshold_* was defined as the contrast level at 80% accuracy, and then the contrast sensitivity was defined as *S* = 1/*C_threshold_*.

#### FMRI experiment

Stimuli were presented with an MRI safe projector (1024 × 768 pixels @ 60 Hz) on a translucent screen placed behind the head coil. Participants viewed the stimuli through a mirror mounted inside the head coil at a viewing distance of 69 cm from the screen. Visual stimuli were identical as those in the psychophysical experiment except that the luminance of the bars and the uniform background were *L_bar_* = 87.5 cd/m^2^ and *L_0_* = 0.4 cd/m^2^, respectively. The purpose of using a dark background was to enhance activations to the texture stimuli. Texture stimuli with different orientation contrast with the background bars (θ = 90°, 15°, or 0°) were presented in separated 30-s blocks, interleaved with 15-second fixation periods (Figure 2C). Each run lasting 270 s, with 6 blocks of stimuli per run (2 blocks for each θ). In each block, there were 60 texture stimuli, each lasting 200 ms followed by 200/300/400 ms of random intervals. The foreground bars were randomly presented in the lower-left or lower-right visual field. Subjects were instructed to maintain fixation and count the number of luminance changes in the fixation cross. Feedback about the numbers of fixation changes was given at the end of each run. A total of 9 runs were collected for each participant. The order of stimulus conditions was balanced across runs and participants.

Two localizer runs were used for each participant to localize the retinotopic regions of interest (ROIs) for the foreground bars. Each run consisted of eight 30-s stimulus blocks, interleaved with 15-s fixation intervals (Figure S2). In each stimulus block, colored naturalistic stimuli (same size as the foreground in the main experiment) were presented at 4 images per second (116.7 ms presentation interleaved with 133.3 ms fixation), at one of the foreground locations. Participants were instructed to maintain fixation and to count fixation changes as in the main experiment.

#### Eye tracking experiment

Eye movements were recorded using an SR Research Eyelink 1000 Plus eye tracker. The centroids of the left eye’s pupil were sampled at 1000 Hz. Visual stimuli were presented on a 27 inches LCD monitor at 1920 × 1080 resolution and 60 Hz refresh rate, at a viewing distance of 100 cm. Participants stabilized their heads using a chin rest. The texture stimuli and fixation task were identical to those employed in the fMRI experiment. Eye gaze positions were preprocessed in the following steps: eye blinks removal (200 ms before and after the eye blinks) with linear interpolation of removed data, linear detrend, baseline correction per stimulus block. A 95% bivariate contour ellipse area (BCEA) was used to quantify the distribution of fixation samples (Steinman, 1965).

### MRI data acquisition

MRI data were collected on a 7T scanner (Siemens MAGNETOM) with a 32-channel receive single-channel transmit head coil (Nova Medical) in the Beijing MRI Center for Brain Research. The gradient coil had a maximum amplitude of 70 mT/m, 200 us minimum gradient rise time, and 200 T/m/s maximum slew rate. Participants used a bite bar to reduce head motion. Functional data of blood nulled and not-nulled images were acquired using a SS-SI-VASO sequence with 3D-EPI readout (Huber et al., 2014; Poser et al., 2010) (0.82-mm isotropic voxels, 26 oblique-coronal slices, FOV = 133 × 177 mm^2^, paired-TR = 5020 ms, TE = 25 ms, TI1 = 1744 ms, FA = 26°, bandwidth = 1064 Hz/pixel, Partial Fourier = 6/8 with 8 POCS iterations, GRAPPA = 3 with FLASH reference). The 3D slab of fMRI acquisition was placed in oblique-coronal orientation (Figure S1C). Images with reversed phase and read directions were acquired for susceptibility distortion correction. T1-weighted anatomical images were acquired using an MP2RAGE sequence (Marques et al., 2010) (0.7-mm isotropic voxels, FOV = 224 × 224 mm^2^, 256 sagittal slices, TE = 3.05 ms, TR = 4000 ms, TI1 = 750 ms, FA = 4°, TI2 = 2500 ms, FA = 5°, bandwidth = 240 Hz/pixel, phase and slice partial Fourier = 7/8, GRAPPA = 3).

#### MRI data analysis

### Preprocessing of functional data

The preprocessing of fMRI data was performed using AFNI (Cox, 1996), LAYNII (Huber et al., 2021) and custom Python code. The following steps were applied for blood nulled and not-nulled volumes separately: EPI image distortion correction with blip-up/down non-linear warping method, rigid-body correction of head motion, spatially up-sampled by a factor of 2 (3dResample in AFNI) (J. Wang et al., 2022). Since the blood nulled and not-nulled volumes were acquired alternately, the 3D timeseries were up-sampled by a factor of 2 (3dUpsample in AFNI) and shifted by TR/2 for BOLD correction. The VASO or CBV-weighted volume was calculated by dividing the nulled by not-nulled volumes (LN_BOCO in LAYNII). To minimize image blur, all spatial transformations were combined and applied to the functional images in one interpolation (3dAllineate in AFNI with sinc method). After per run scaling as percent signal change, a general linear model (GLM) with a canonical HRF (BLOCK4 in AFNI) was used to estimate the VASO and BOLD signal change from baseline. The fMRI signals in all cortical depths were fitted equally well by convolving the stimulus regressor with the canonical HRF (Figure S9). Thus, the GLM analysis should not introduce bias across cortical depth. Head motions and low-order drifts were included as regressors of no interest in the GLM.

### Surface segmentation

The T1-w anatomical volume was aligned to the mean EPI image after motion correction, and segmented into white matter (WM), gray matter (GM), and cerebrospinal fluid (CSF) compartments using FreeSurfer with the “high-res” option (Fischl, 2012). FreeSurfer’s segmentation results were visually inspected and manually edited to remove dura mater and sinus, etc., ensuring accurate surface segmentation (Figure S1A). High-density surface mesh was generated by a factor of 4 to better align with the up-sampled volume grid (Polimeni et al., 2018). The cortical depth profiles were constructed using the equi-volume model (Waehnert et al., 2014). Two equi-volume intermediate surfaces between the WM and pial surfaces were generated using mripy (https://github.com/herrlich10/mripy), dividing the GM into three equi-volume layer compartments (Figure S1A). For each voxel, the layer weight (volume percentage among WM, CSF and the three layers compartments) were calculated and subsequently used in a spatial regression approach to unmix layer activity (Kok et al., 2016).

### ROI definition

Anatomical ROIs of the early visual areas (V1-V2) were defined on the cortical surface by a 7T retinotopic atlas of the Human Connectome Project using Neuropythy tools (Benson et al., 2014, 2018). The anatomical ROI for IPS was taken from the Wang15 atlas (L. Wang et al., 2015). Vertices with positive activation from baseline in the localizer (p < 0.05, uncorrected) were selected within the anatomical ROI as the foreground ROI (Figure 2D). Accordingly, surround ROIs in V1 and V2 were acquired by subtracting the foreground ROI from vertices with significant activation to the texture stimuli (p < 0.05, uncorrected). Surface ROIs were then projected to the EPI volume to select voxels in a column-wise manner.

### Visual Field Map Reconstruction

The visual field map of saliency-related signals was reconstructed for each cortical depth using the saliency-dependent (θ_90_ - θ_15_) CBV responses and the population receptive field (pRF) maps from the Benson atlas (Benson et al., 2018; Dumoulin & Wandell, 2008). To account for individual differences in the pRF maps, the pRF locations were linearly transformed by aligning the peak of localizer activation to the foreground location. The visual field map was reconstructed by multiplying the each node’s CBV response with its pRF map and then summed across all nodes (Mo et al., 2018). Finally, the reconstructed map was normalized (divided) by the maximum response across all cortical depths, and then averaged across participants.

### Layer-dependent effective connectivity

A generalized psychophysiological interaction (gPPI) method was used to investigate the effective connectivity across cortical depths in V1 and IPS (McLaren et al., 2012). CBV timeseries of the foreground ROI were averaged within each cortical depth of V1 and IPS. Due to the uncertainty about CBV HRF, the PPI term was calculated by the dot product of the stimulus regressor (boxcar function, shifted by one TR due to the HRF delay) and the seed timecourse without deconvolution (O’Reilly et al., 2012; Sharoh et al., 2019). The GLM includes the seed region time course, stimulus regressors, PPI terms of the three θ conditions, along with baseline regressors and head motion parameters. Saliency-dependent functional connectivity was defined as the difference between the PPI beta weights of the θ_90_ and θ_15_ conditions.

### Statistical analysis

Two-way repeated-measures (rm) ANOVA with cortical depths (deep, middle, and superficial) and θ (90°, 15°, and 0°) as within-subject factors were performed on the ROI-averaged CBV responses. All data met the sphericity assumption of ANOVA. In cases where the sphericity assumption was violated, p values were adjusted using the Greenhouse-Geisser correction. One-way rm ANOVA was performed to evaluate the difference in saliency-related signals across cortical depths, followed paired t-tests if there was a significant difference. No FWE correction is needed for paired t-tests followed by a significant one-way ANOVA with three levels (Levin et al., 1994).

Pearson’s r was calculated to assess the correlation between the fMRI saliency sensitivity *SS_fMRI_* and the behavioral saliency sensitivity *SS_behavior_*. Data were assessed for multivariate normality using Shapiro-Wilk test (all p > 0.05). A permutation test was performed to correct for the family-wise error (FWE) across ROIs. In each permutation, Pearson’s r was calculated after shuffling the correspondence between fMRI signal and perceptual sensitivity across participants. The largest r value was taken and the permutation was repeated to generate a null distribution, from which the FWE was derived for the original r values.

## Acknowledgement

This study was supported by STI2030-Major Projects (2022ZD0211900 and 2021ZD0204200 to P.Z.), National Natural Science Foundation of China (31871107 and 31930053 to P.Z.). Hunan Provincial Natural Science Foundation (2024JJ6313 to C.W.L), Scientific Research Foundation of Hunan Provincial Education Bureau (24B0058 to C.W.L). NIH Intramural Program of NIMH/NINDS (#ZIC MH002884 to L.H.).

## Supplementary figures

**Figure S1.**
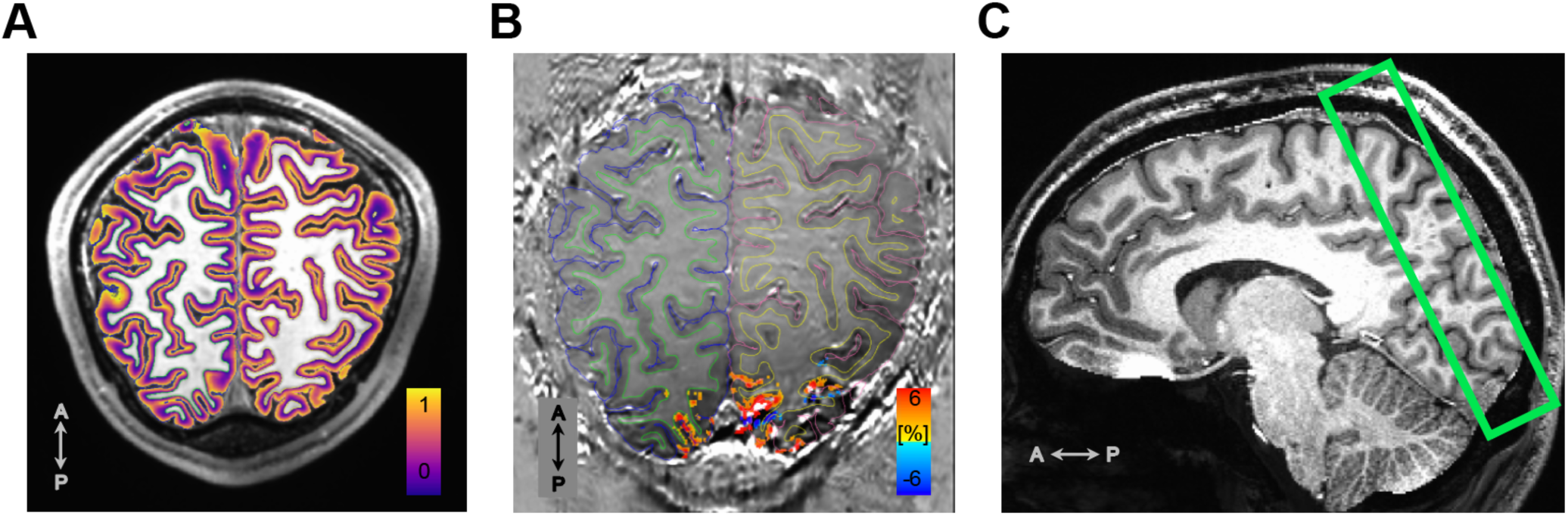
(A) Normalized cortical depth map overlayed on the T1w anatomical image in a representative participant. The equi-volume depth at 0 and 1 correspond to the WM and Pial surfaces, respectively. (B) VASO activations (90°+15°+0°, p < 0.001 uncorrected) overlayed on the mean VASO image. Green and yellow lines indicate the WM surface, while the blue and pink lines denote the pial surface. (C) The green box indicates the 3D slab of VASO fMRI acquisition.

**Figure S2.**
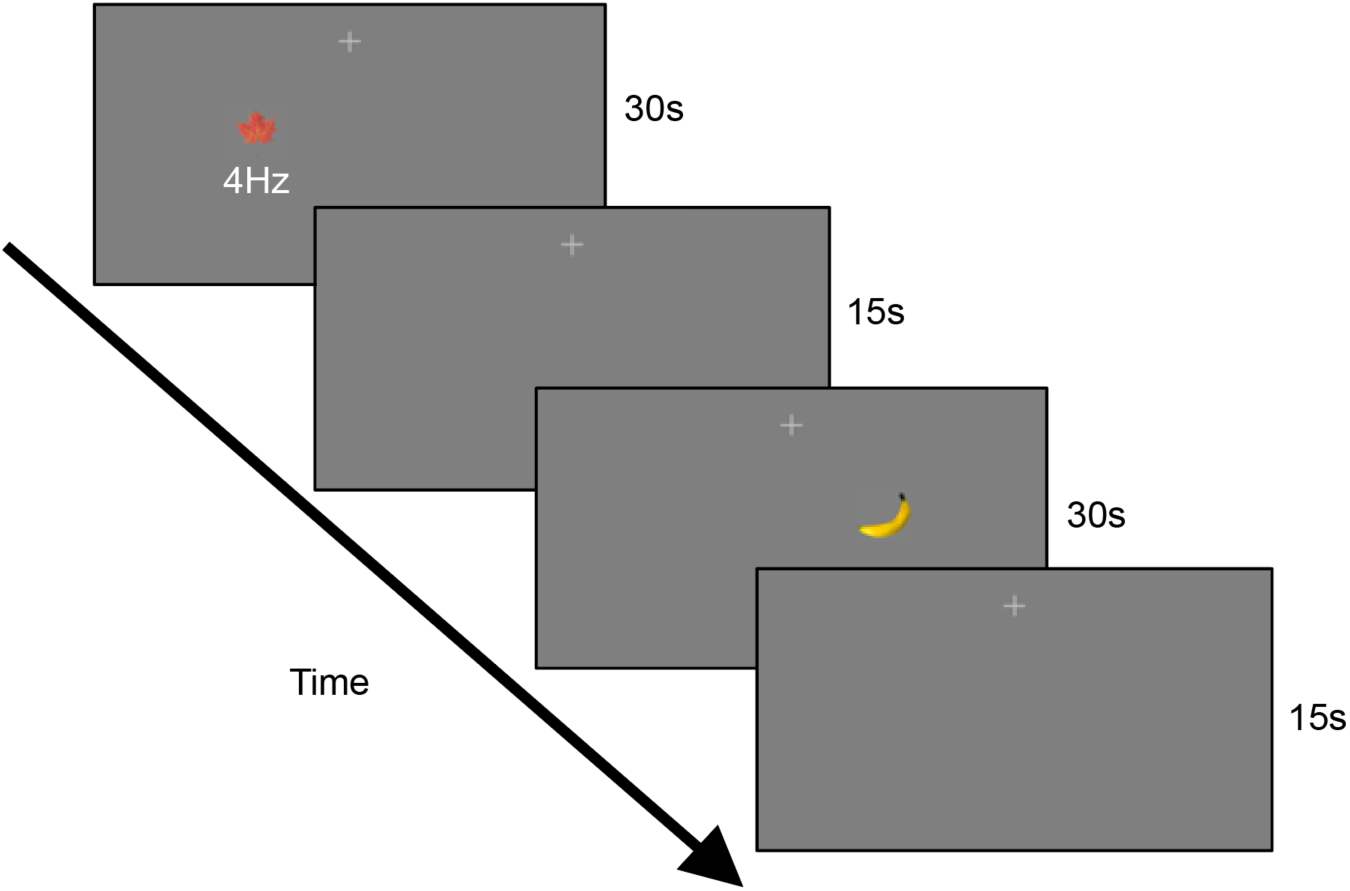
Stimulus and procedure of the localizer runs. Naturalistic stimuli were presented at 4 images per second in the lower-left or lower-right quadrants in separate stimulus blocks, interleaved with 15-second fixation periods. The size and location of localizer stimuli matched the foreground region in the main experiment.

**Figure S3.**
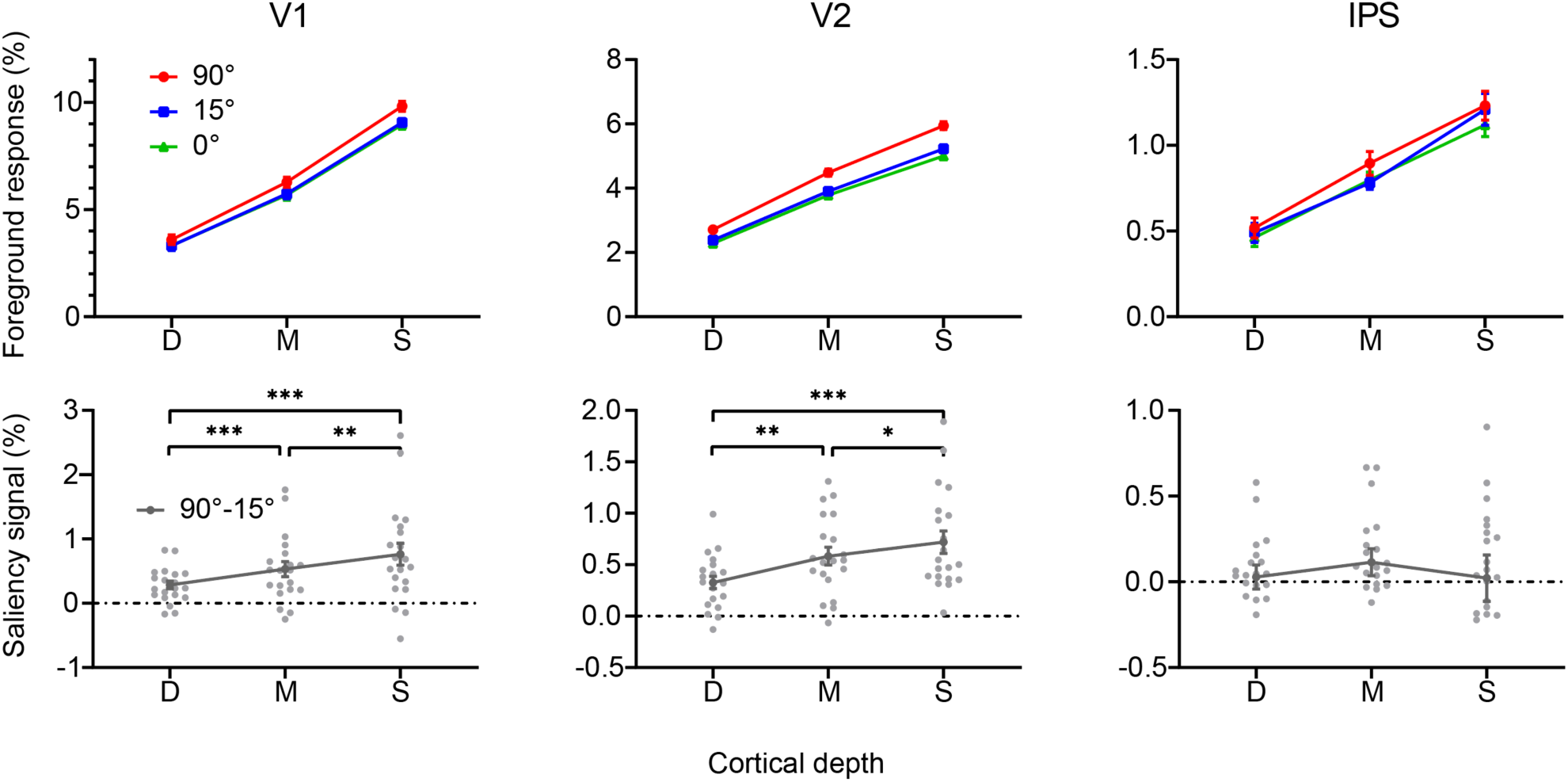
Normalized BOLD responses in the foreground ROIs of V1, V2, and IPS. Top panel: BOLD response in different depths of V1, V2, and IPS in 90°, 15°, and 0° orientation contrast conditions; Bottom panel: Calculated from top panel, the response difference between 90° and 0° foregrounds. Error bars represent the standard deviation of the mean. *, ** and *** indicate p < 0.05, p < 0.01, p < 0.001.

**Figure S4.**
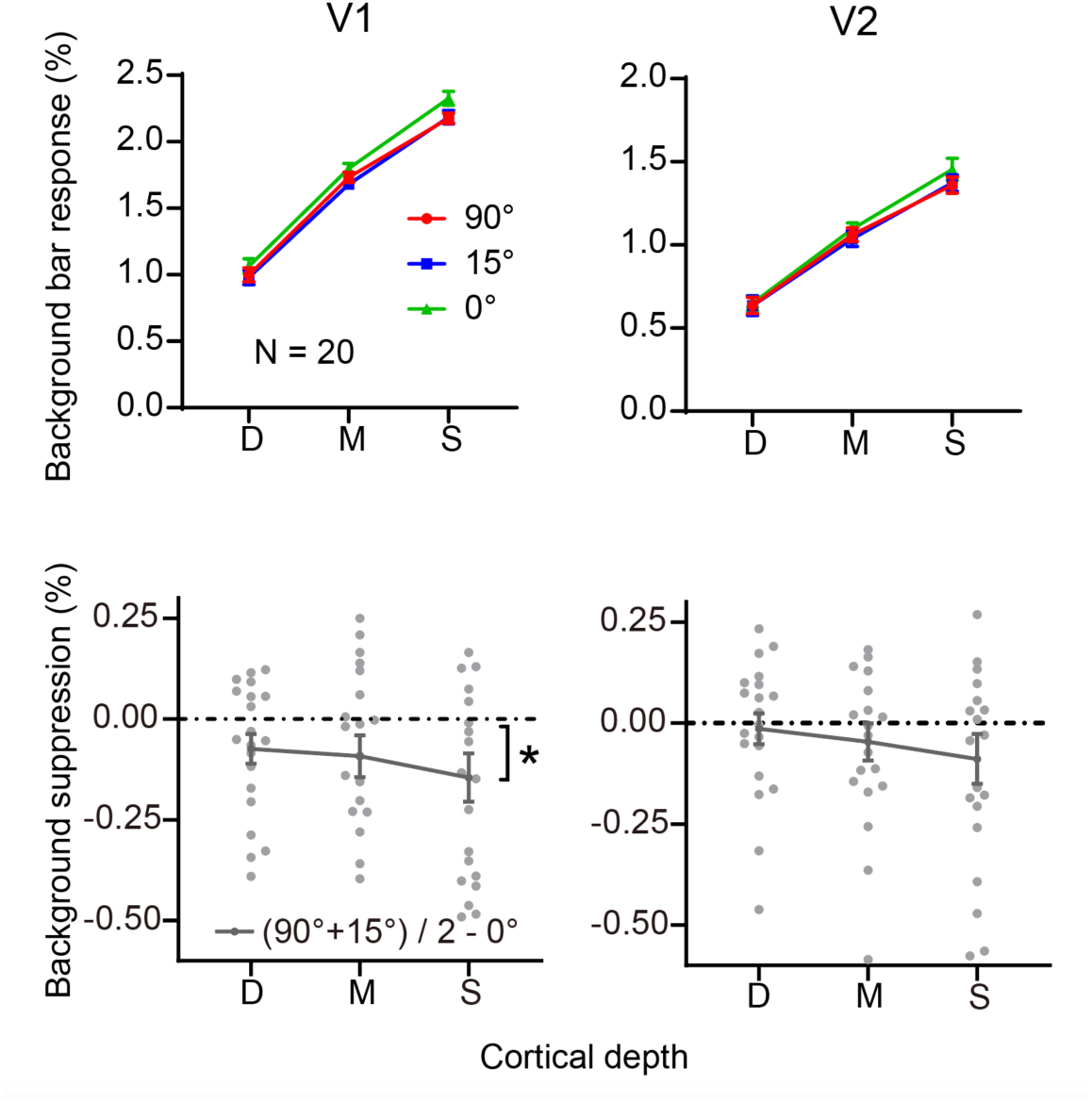
Upper: normalized CBV responses in the background ROIs of V1 and V2. A significant effect of orientation contrast (θ = 90°, 15°, and 0°) was found in V1 (F_2, 38_ = 3.700, p = 0.034) and a similar trend in V2, suggesting weaker background activity in the 90° and 15° conditions compared to the 0° or the uniform texture condition. No significant difference was found between the two θ conditions (F_1, 19_ = 0.365, p = 0.553, BF_10_ = 5.601×10^-11^). Lower: The suppression effect was calculated as the response difference between the mean of 90° and 15° conditions and the 0° condition ((90°+15°)/2-0°). A significant suppression effect was found only in the superficial depth of V1 (t19 = -2.877, p = 0.023, Holm corrected across cortical depths). These results suggest a suppression effect of background activity in the superficial layers of V1, independent with the orientation contrast between the foreground and the background bars. Each gray dot represents one participant. Error bars indicate SEM. * indicates p < 0.05. D, M, S indicate deep, middle, and superficial depth, respectively.

**Figure S5.**
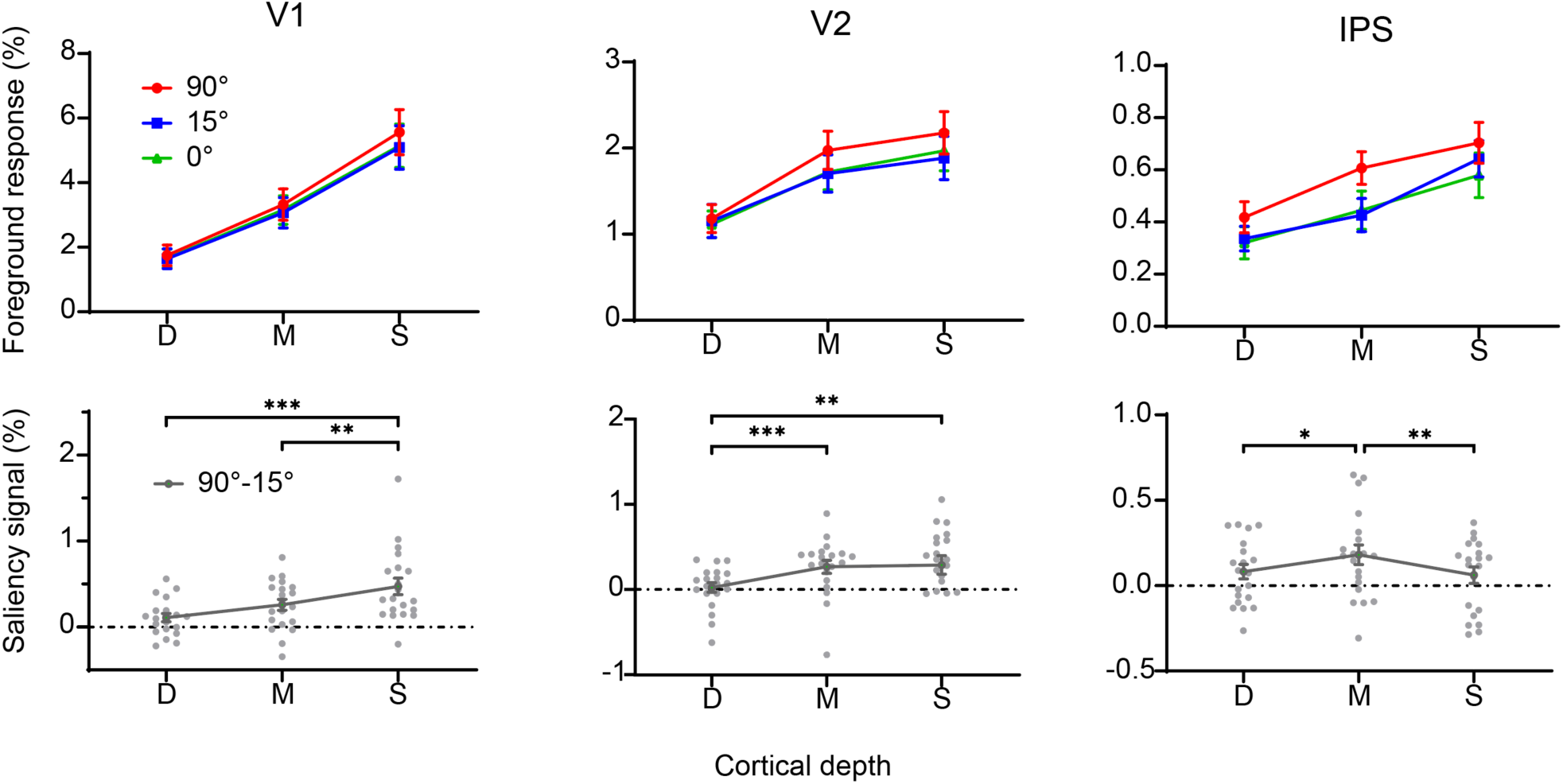
Unnormalized (or original) CBV responses in the foreground ROIs. Conventions are identical as in Figure 3A.

**Figure S6.**
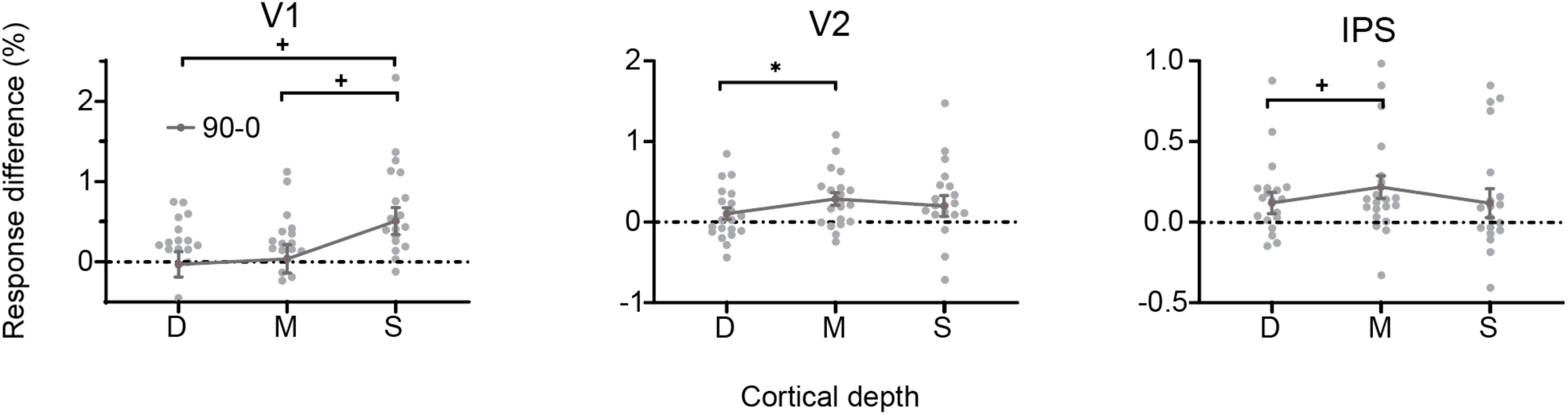
Normalized CBV response difference in the foreground ROIs between 90° and 0° orientation contrast conditions. Similar laminar profile with Figure 3A bottom panel. Error bars represent the standard error of the mean. * p < 0.05, + p < 0.1.

**Figure S7.**
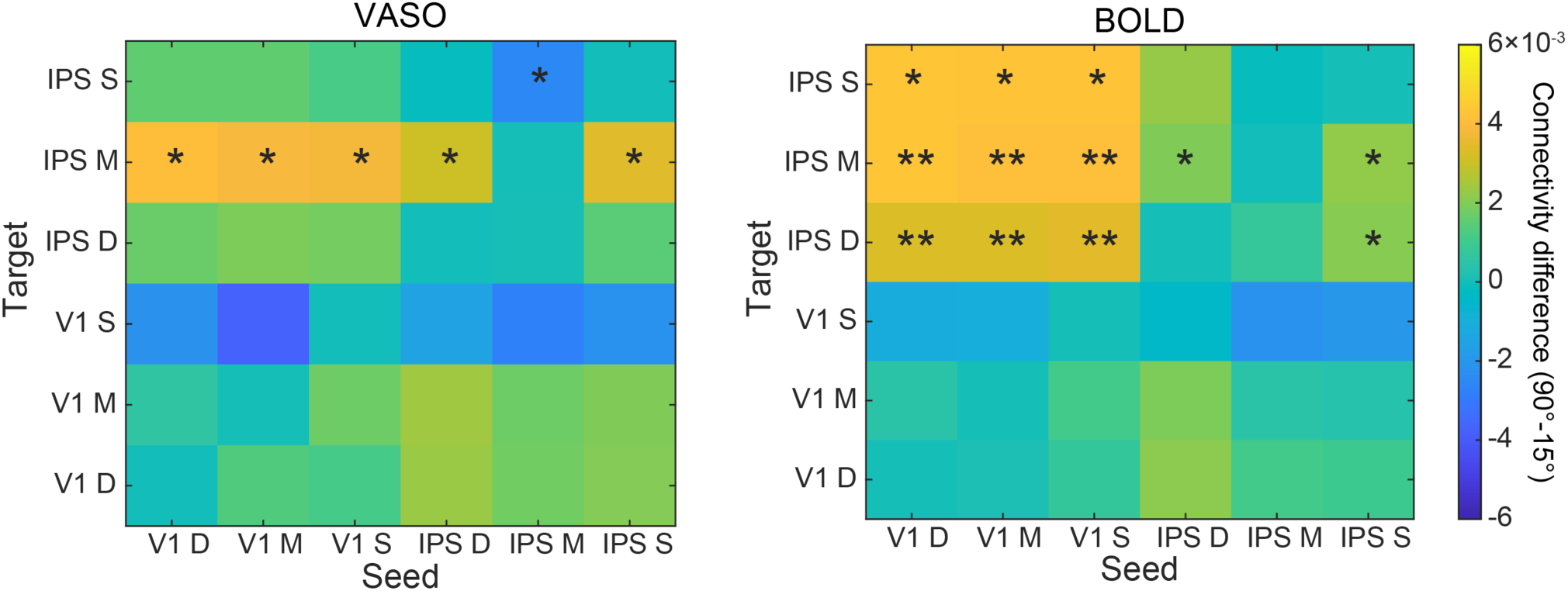
The gPPI connectivity matrix across three cortical depths in V1 and IPS. Columns and rows correspond to the seed and target ROIs, respectively. The color scale indicates the beta difference of interaction terms (beta(90°)-beta(15°)). * p < 0.05, ** p < 0.01, uncorrected.

**Figure S8.**
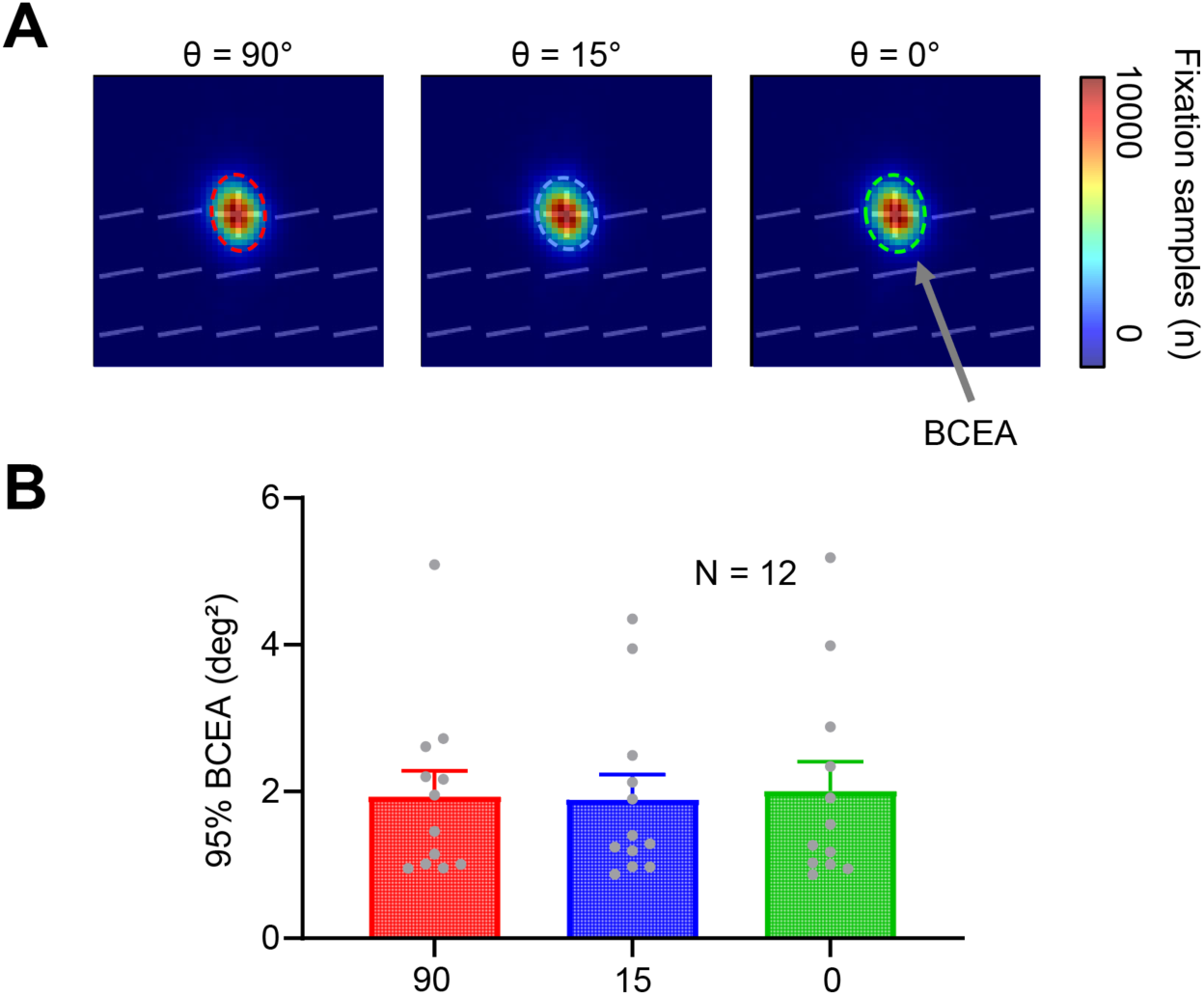
(A) The group-averaged heat maps of fixation distribution in the entire session. (B) The bivariate contour ellipse area (BCEA) of fixation distribution showed no significant difference across θ conditions (F_2,22_ = 0.291, p = 0.750, BF_10_ = 0.227). Error bars represent the standard deviation of the means.

**Figure S9.**
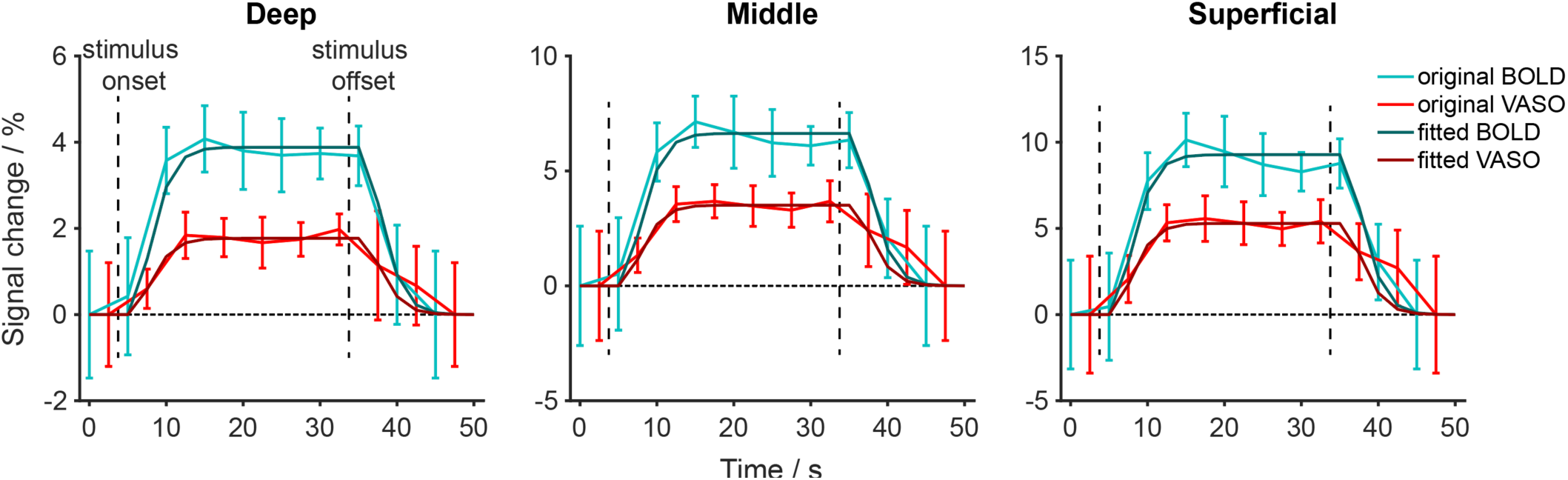
VASO and BOLD response timecourses in deep, middle, and superficial ROIs of the foreground in V1, and the fitted response with GLM using a canonical HRF (BLOCK4 in AFNI). Error bars represent SEM across participants.

